# Spread of *Xanthomonas vasicola* pv. *musacearum* within banana mats: implications for Xanthomonas wilt management

**DOI:** 10.1101/2021.11.04.467225

**Authors:** Walter Ocimati, Anthony Fredrick Tazuba, Guy Blomme

**Affiliations:** Alliance of Bioversity International and CIAT, P.O. Box 24384, Kampala, Uganda; Alliance of Bioversity International and CIAT, c/o ILRI, P.O. Box 5689, Addis Ababa, Ethiopia

**Keywords:** bacteria, corm, inoculate, incubation period, incidence, suckers

## Abstract

Xanthomonas wilt (XW) of banana caused by *Xanthomonas vasicola* pv. *musacearum* (Xvm) does not spread to all plants physically interconnected through the rhizome when one or a few are diseased. However, the factors behind this incomplete systemic spread of Xvm are not fully known yet could inform XW management. This study explored the effect of Xvm inoculum amounts, number and size of suckers, sucker positioning on mother plant corms and other mother plant corm attributes on sucker colonization. A shorter (p <0.05) incubation period (17.9 vs 21.1 days) and higher (p<.001) cumulative number of symptomatic leaves (5.2 vs 1.6 leaves) was observed when all (high inoculum) compared to two leaves (low inoculum) were inoculated. Xvm was recovered in corms at 29 days post inoculation (dpi) in both treatments with no differences (p >0.05) in proportions of corms with Xvm between the treatments. However, Xvm was recovered earlier and at a higher frequency in suckers when all leaves were inoculated. Lower Xvm recoveries occurred in the lower corm sections to which most suckers were attached relative to the middle and upper corm sections. Xvm incidence in corms increased with the number of attached maiden suckers, and the dpi while it declined with increasing mother plant and corm height. Thus, Xvm spread within mats is influenced by the amount of inoculum and the physiological stage (e.g., height) of the plant and attached suckers. The position of suckers, predominantly at the bottom of corms also protects them from infection. Measures that reduce Xvm inoculum build-up in mats are thus crucial for minimizing within mat XW spread.

## Introduction

Xanthomonas wilt (XW) of banana and enset caused by the bacteria *Xanthomonas vasicola* pv. *musacearum* (Xvm) is an important disease of banana in the East and Central African region. First observed on enset in the 1930’s in Ethiopia (Castellani 1939), XW was subsequently observed in that same country on banana in 1974 (Yirgou and Bradbury 1974). Outside Ethiopia, XW was first observed in 2001 in Uganda (Tushemereirwe et al. 2003) and the Democratic Republic of Congo (Ndungo et al. 2006) and has since spread to Rwanda (Reeder et al. 2007), Burundi, Tanzania and Kenya (Carter et al. 2010).

Knowledge of the epidemiology of XW disease has been crucial in the design of its control strategies. For example, studies to understand the within plant and mat spread of Xvm (Ocimati et al. 2013a, 2015; Blomme et al. 2017a; Ntamwira et al. 2019) have shown that Xvm does not spread to all physically attached plants in a mat when one or a few plants are visibly diseased, a phenomenon referred to as “incomplete systemic” spread of Xvm. This phenomenon explains the success of the single diseased stem removal (SDSR) technique in XW management (Blomme et al. 2017a, 2019). However, the factors behind the incomplete systemic spread of Xvm in a mat are not fully understood.

Several factors have been reported or postulated to account for this incomplete systemic spread of Xvm. For example, the removal of a mature infected plant with early male bud wilting symptoms has been reported to potentially prevent Xvm from reaching the corm of the infected plant and hence other physically attached lateral shoots (Ssekiwoko et al. 2010; Ocimati et al. 2013a, 2015). However, early male bud wilting symptoms are not often easily noticed by farmers whereas infections could also be introduced through the other vegetative parts such as the leaves. At higher XW symptom severity levels such as yellowing of leaves, pre-mature fruit ripening and death of infected plants, Xvm has been reported to be already present in the corm tissues (Ssekiwoko et al. 2010; Ocimati et al. 2013a, 2013b). However, field recoveries have also been observed in heavily diseased fields (with plant incidence levels as high as 80%) where infected plants often exhibit more severe XW symptoms (Blomme et al. 2017a, 2019; Ntamwira et al. 2019).

Ocimati et al. (2013b) observed a markedly lower XW incidence and higher incubation period for plants inoculated at the corm level through de-suckering compared with plants inoculated through de-leafing. These authors thus suggested a possible delay in Xvm colonisation of corm tissues due to its compact nature. In a more recent study, the corm of the resistant enset cultivar ‘Mazia’ was however observed to be significantly softer than that of the susceptible enset cultivars ‘Arkiya’ and ‘Kelisa’ (Said et al. 2020), suggesting that corm hardness may not necessarily be responsible for the observed incomplete systemic spread of Xvm.

It was also theorised that incomplete systemicity could be due to the reduced dependence of more mature maiden suckers on parent plants (maiden suckers have their own fully developed root systems and mature leaf canopy and were hence postulated to depend less on the mother plant for water and nutrients), thus reducing the likelihood of them becoming infected via the mother plant. Ntamwira et al. (2019) however observed higher XW infections in bigger suckers (maiden suckers) compared to smaller ones (peepers and sword suckers) attached to infected mother plants, challenging the hypothesis that bigger suckers are more self-reliant and less susceptible to infections from the attached mother plants. Ntamwira et al. (2019) however observed a reduction in incidence of XW infections in attached suckers when artificially inoculated mother plants were timely removed, an act that possibly prevented the build-up of XW inoculum in the mother plant and wider mat. This study suggested that the amount of Xvm inoculum in an infected plant potentially influences XW infections in other physically attached plantlets.

Suppression of Xvm by endophytes at the corm level had also been postulated to explain incomplete systemic spread of Xvm at the mat level (Karamura et al. 2016; Blomme et al. 2017a). Studies on beneficial micro-organisms carried out by Were (2016) showed promising levels of Xvm suppression by bacterial isolates from banana tissues in *in vitro* studies, while Abayneh (2010) reported a 56 to 75% reduction in XW incidence in pot experiments for plants inoculated with beneficial endophytes through leaf and pseudostem tissues. Studies to validate these findings in the field are however still lacking.

This study built on the above studies through exploring the effect of i) different Xvm inoculum amounts (through inoculating different numbers of leaves on a mother plant) and ii) the location of suckers on the mother plant corm tissue (sucker closeness to point of attachment on corm of infected leaves) on within mat Xvm, specifically on the colonisation of the attached suckers/ lateral shoots. It is hypothesized that Xvm spread from the mother plant to the physically attached suckers could be influenced by a) the amount of Xvm inoculum in the mother plant or plant through which the infection is introduced, b) the position of a sucker on the mother plant corm relative to the inoculated leaf (i.e., the closer a sucker is attached to the insertion point of the inoculated leaf/ leaf sheaths of the mother plant, the higher the chance the sucker gets infected). This knowledge is anticipated to help in fine-tuning the XW management through the current cultural practices.

## Materials and methods

### Field experimental set up

The field experiment was conducted in an isolated site within Kifu forest (00°280N, 32°440E), in Mukono district, central Uganda. Kifu has a mean daily temperature of 25°C, and a mean annual rainfall of 1100 mm that is bimodally distributed (March–May and September–November). A total of 100 East African highland banana cultivar “Mbwazirume’ (*Musa* AAA genome group) suckers were planted in March 2019 at a spacing of 2 × 2 m. The fields were established using corms of maiden suckers obtained from a XW-free field. Effort was made to obtain corms of approximately the same size (~2.6 kg; 45.3 cm circumference; 29.5 cm from top to bottom of corm). From the sixth month after planting, mother plants (70 plants) with at least two sword suckers (i.e., lateral shoots with lanceolate leaves) were artificially inoculated with a suspension of Xvm.

### Bacteria inoculum preparation

The Xvm inoculum was obtained from two plants with characteristic XW symptoms located in a single experimental field at Kifu in Mukono district, Uganda. Plants from a single field were used to minimise variation in virulence of the isolates used. The two plants were cut down with a sterile machete, the pseudostems cut transversally into smaller 30 cm length portions and allowed to ooze bacteria for about 30 to 45 min. The yellow ooze, characteristic of Xvm was then scrapped off the surface of the pseudostem sections into a 50 mL falcon tube. A bacterial suspension was then prepared by thoroughly mixing 50 mL of the ooze with 950 mL of double-distilled water in a sterile 1000 mL conical flask.

### Inoculation treatments

For half of the mother plants with at least two sword suckers (i.e., 35 plants or mats), all the functional leaves were inoculated [at petiole level] to ensure a higher Xvm inoculum load and increase the chances for a more uniform Xvm colonization of the corm. For half, only 2 functional leaves (youngest and fully open leaves) located on the same side of the mother plant were inoculated. Treatments were randomly assigned. To inoculate the plants, 1 mL of Xvm suspension was injected into the middle section of the leaf petiole about 10 cm from the pseudostem of each inoculated leaf. The bacterial suspension was thoroughly stirred each time a new suspension was sucked into the syringe to inject a more or less equal number of bacteria in each leaf petiole. To introduce the bacteria, the needle was inserted at a 30–45-degree angle into the leaf petiole and Xvm suspension gently injected into the petiole. Ribbons where then attached onto the inoculated leaves to mark/distinguish them.

### Data collection

At inoculation, the following mother plant growth traits were measured: pseudostem circumference at soil level, plant height and number of functional leaves. In addition, the number of suckers attached to the mother plants and their respective heights were assessed (Table 1).

**Table 1.**
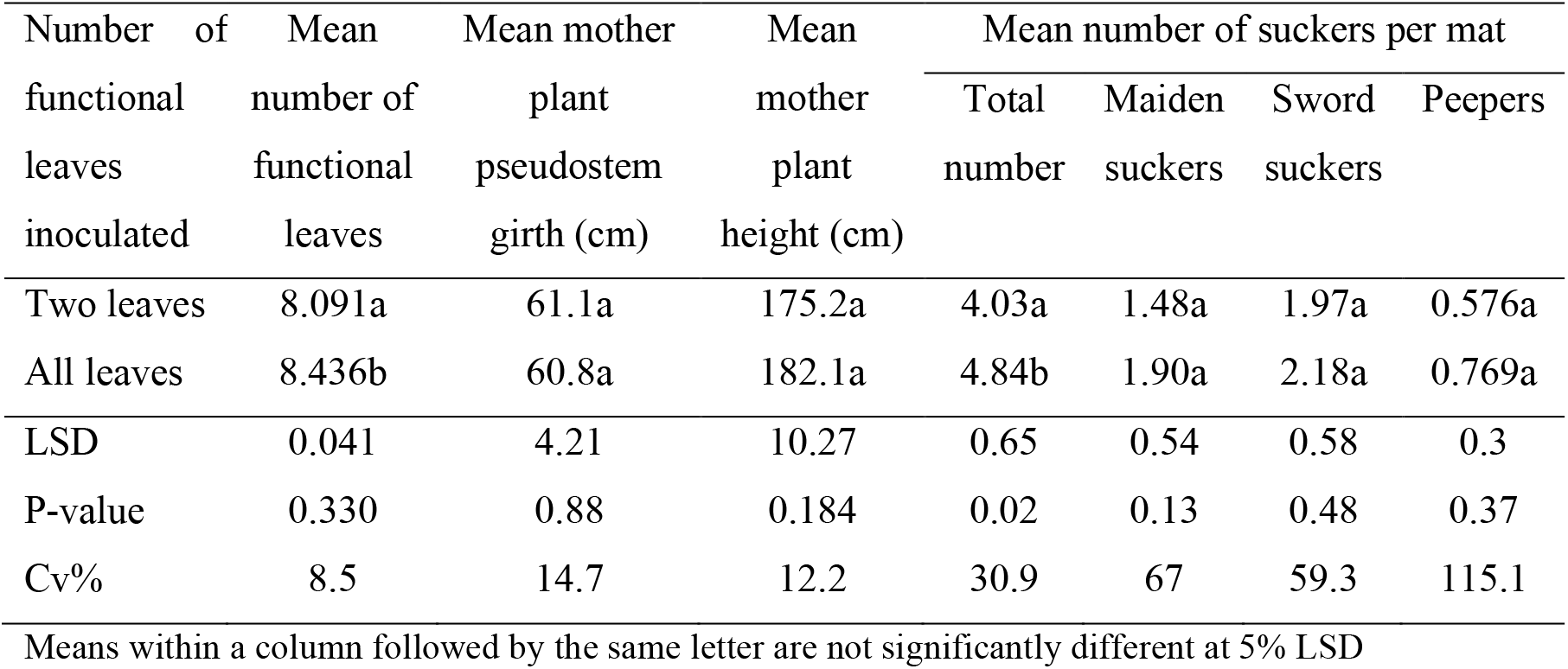
The mean number of functional leaves, pseudostem girth at soil level and height of mother plants; mean total number of suckers; and the mean number of maiden suckers, sword suckers and peepers per mat at time of mother plant inoculation with *Xanthomonas vasicola* pv. *musacearum*.

Except for the total number of suckers attached to the mother plants that differed significantly (P<0.05), no significant (P>0.05) differences were observed in the other variables between the ‘all leaves’ and ‘2 leaves’ treatment plants (Table 1). This suggests that the attributes of the plants at inoculation may not have profoundly affected the outcome of the study. About half of the inoculated plants i.e., 17 mats of each category of inoculation treatment (i.e., all leaves vs 2 leaves) were randomly assigned to be observed for symptom development in the mother plant and attached suckers. Data collected on these plants included the time to first symptom development in the mother plant and suckers (i.e., XW incubation period), number of symptomatic leaves, time to symptom expression for each visibly diseased leaf and the number of suckers that developed disease symptoms.

For the remaining mats (36 mats), i.e., 18 mats for either inoculation treatment, 3 mats each were randomly sampled at 2, 4, 6, 8, 10 and 12 weeks after inoculation for enumeration of Xvm in the laboratory. Entire mats comprising the mother plant and attached sucker corms with the roots (Fig. 1A) were dug out using hoes sterilized in between plants.

**Figure 1.**
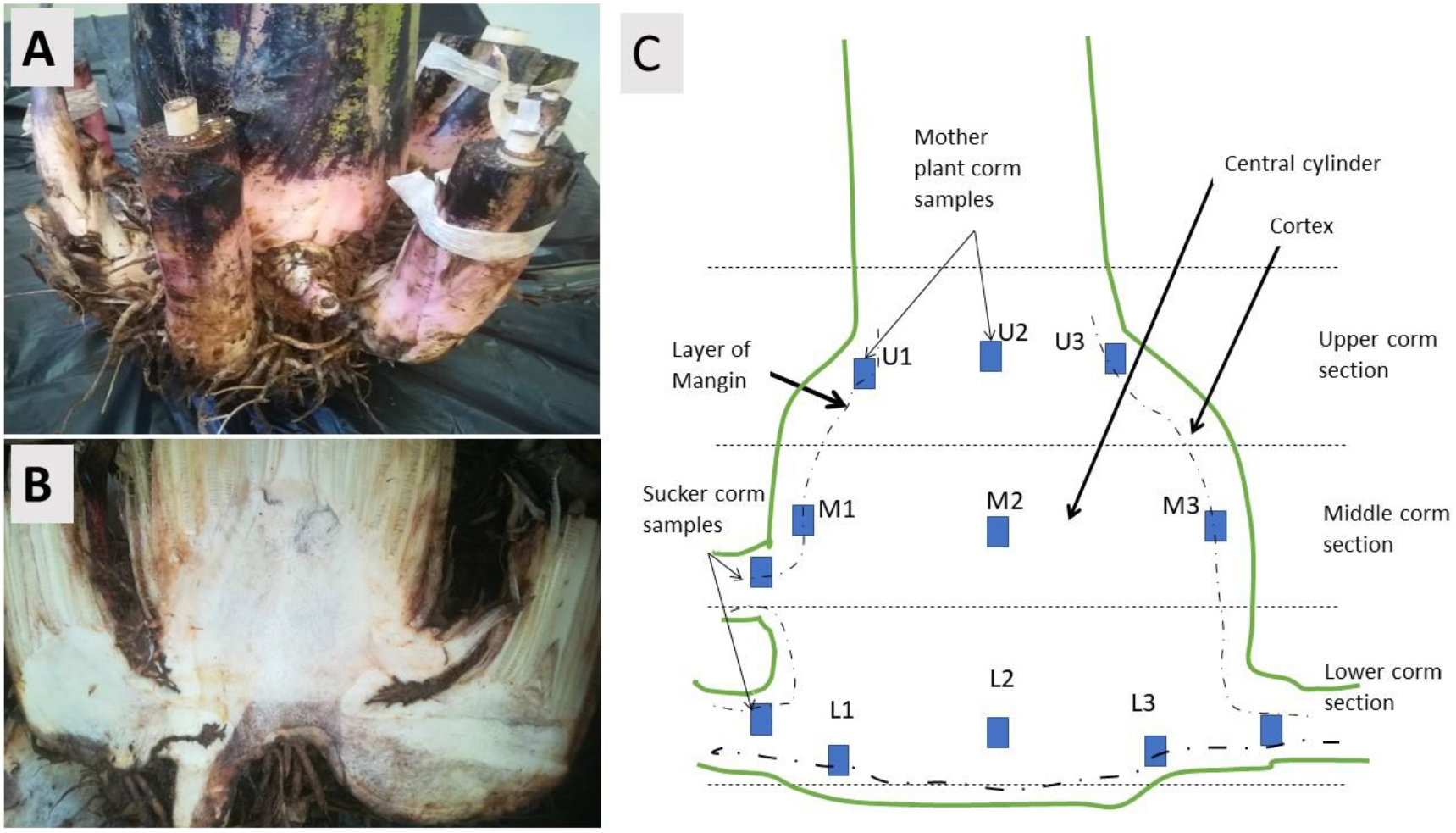
Mat comprising of the mother plant and physically interconnected lateral shoots or suckers (A); a longitudinally split corm revealing different sections of the mother corm and the sucker corms (B) and an illustration of the longitudinal section of a split corm showing points on the corms from where samples were extracted for laboratory analysis (C). In (C), cortex is the zone outside the layer of Mangin while the central cylinder is the zone of the corm inside the layer of Mangin.

The mother plant roots were sampled and put into separate paper bags, attached suckers labelled, and the roots directly attached to the suckers aseptically and separately sampled into separate collection bags. In the field, pseudostem samples were also aseptically cut from the main shoot and the attached suckers at 5 cm above the corm tissues for Xvm isolation in the laboratory. As much as possible, soil attached to the interconnected corms of the mother plant and suckers was carefully removed in the field. The entire cluster of corms was subsequently transported to the laboratory where it was thoroughly washed to remove the remaining soil debris.

In the laboratory, the corms were longitudinally split/dissected using aseptic tools (knives and machetes) along the insertion points of the suckers exposing the different layers (cortex, layer of Mangin (cambium ring in the corm), central cylinder) of both the mother plant and attached sucker corms (Fig. 1B). As many dissections of the mother plant corm were carried out as there were tagged suckers for sampling in the lab. The distance from the point of insertion of the sucker on the mother plant corm to the mother plants’ apical meristem was measured along the outer corm surface of the mother plant corm and on the longitudinal section using a direct line from the apical meristem to the point of insertion of the sucker. The height of the corm from apical meristem to the bottom of the corm was also measured. The surface corm tissue of the longitudinal sections was then sterilized with a 3.5% (v/v) sodium hypochlorite (NaOCl) solution and 70% (v/v) ethanol to eliminate potential contamination that could have occurred during the longitudinal splitting of the corms. Subsequently, the longitudinal corm surface was thoroughly rinsed with sterile water to remove any excess NaOCl and ethanol.

From the surface of the longitudinally cut corms, at least 12-16 pieces of 3 cm thick cube-shaped corm tissues were cut out using sterile blades at equal distances i) along the cortex-layer of Mangin and central cylinder of the corm; and ii) at insertion points of the sucker corms to the mother plant corm (Fig. 1C). A cross-sectional cut of the pseudostem 10 cm from the sucker corm of the attached suckers was also sampled. The surfaces of the cube-shaped corm tissue samples were peeled off to remove any contaminants and portions previously drenched with NaOCl and ethanol. For the pseudostem tissue, the outer leaf sheaths were wiped with cotton wool soaked with ethanol followed by wiping repeatedly with cotton soaked in sterile distilled water. The corm and pseudostem samples were subsequently macerated in sterile mortars. The macerated tissues where then transferred into 2.0 mL eppendorff tubes, 1 mL of sterile water added, briefly vortexed to dislodge the bacteria. 1.0 mL of the suspension was aliquoted into a 1.5 mL eppendorf tube, serially diluted to 10^−2^. 10 μL of the zero and 10^−2^ dilutions were aseptically spread plated on semi-selective yeast peptone glucose agar petri plates. The plates were then incubated for a period of 3 days and the presence and where feasible the number of characteristic Xvm colonies recorded. The Xvm bacteria were also confirmed using an Xvm-specific primer using a polymerase chain reaction before inoculation into the banana plants and at sample assessment stage (Nakato et al. 2018).

The hardness/ compactness of different sections of three randomly selected vegetative and flowering stage mother plant corms were determined using a soil penetrometer (Scale 0-4.5 Kgf/cm^2^; ELE Pocket Penetrometer 29-3729; https://www.ele.com/product/pocket-penetrometer-) modified to have a sharp pointed (conical shaped iron) tip. Corm tissue hardness was assessed for the upper, middle, and lower corm cortex and central cylinder sections.

### Data analysis

Analysis of variance comparing the disease progression in the above ground parts of the mother plants, Xvm incidence in the mother plant pseudostem and corm parts and sucker tissues between the two treatments was computed using the R statistical package (R Core Team 2018). Paired t-tests were used to determine the relationship between Xvm presence in mother plant tissues and the attached suckers. To determine the most important factor explaining the incidence of Xvm in the different corm sections, a logistic regression of Xvm incidence as a dependent variable against a range of independent variables was conducted using the R statistical package (R Core Team 2018). The independent variables included corm height, distance from point of sucker insertion on mother plant corm to apical meristem along the outer corm surface and the longitudinal section from the apical meristem to the point of insertion of the sucker. Other independent variables included the mother plant height, mother plant pseudostem girth at soil level, number of functional leaves on mother plant, total number of suckers and number of suckers disaggregated into maiden suckers, sword suckers and peepers at inoculation, the days post inoculation and the treatments (i.e., inoculation of 2 or all leaves). The R package and Ms Excel were used to generate visuals.

## Results and discussion

### XW incubation and severity under different inoculation scenarios

It took a significantly (p =0.015) shorter time (mean of 17.9 days) for the first XW characteristic symptoms to appear on plants in which all leaves were inoculated compared to 21.1 days when only two leaves were inoculated (Fig. 2A). These results agree with findings of earlier inoculation studies on vegetative stage banana plants (Ocimati et al. 2013b). Leaves turned yellow and eventually started wilting, typical of a XW infection. The leaves often looked as if scorched by fire and sometimes broke halfway the midrib.

**Figure 2.**
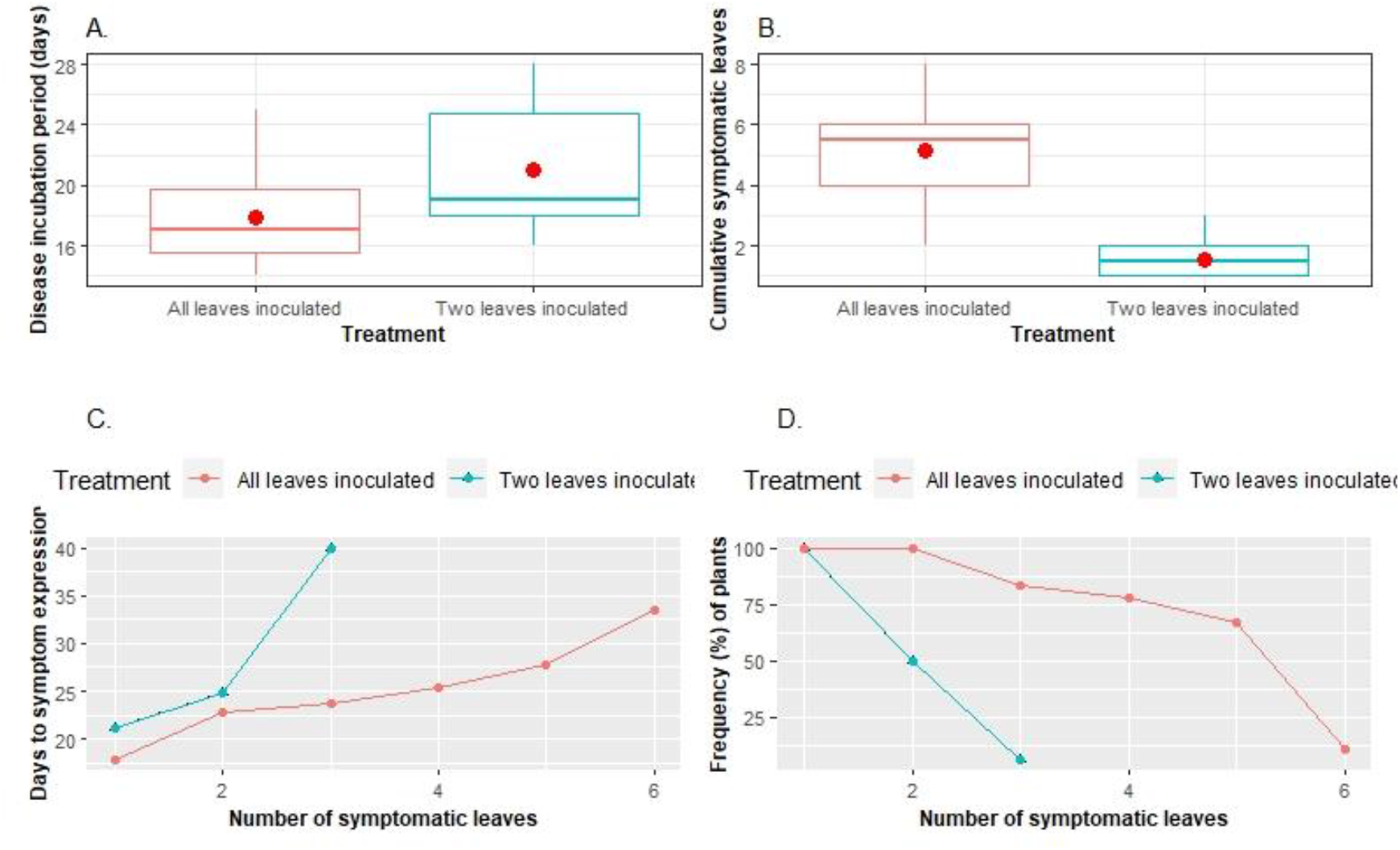
Xanthomonas wilt incubation period (A) and cumulative number of symptomatic banana leaves (B) for mother plants in which either two or all leaves were inoculated; time to symptom expression in different leaves (C) and the percentage frequency of plants with varying numbers of symptomatic leaves (D).

A significantly higher (p <001) mean cumulative number of leaves (5.2 leaves) showed XW symptoms on mother plants in which all leaves were inoculated compared to 1.6 leaves for those on which only 2 leaves were inoculated (Fig. 2B). A maximum of three leaves, with the third leaf showing symptoms 40 days post inoculation (dpi) was observed in mother plants on which two leaves had been inoculated (Fig. 2C). In contrast, up to 6 leaves showed XW symptoms in the treatment in which all leaves had been inoculated, with a lower incubation period of about 32.6 days in the 6^th^ leaf. For both treatments (all and 2 leaves) at least one inoculated leaf showed XW symptoms in 100% of the inoculated plants. Only 50% and 6.3% of the plants in which two leaves had been inoculated showed symptoms in only 2 and 3 leaves, respectively. 100% of the “all leaves inoculated” treatment had 2 symptomatic leaves, the percentage dropping steadily to 11% for 6 symptomatic leaves in a plant (Fig. 2D). These findings suggest that the higher the amount of Xvm inoculum received by plants the higher their susceptibility and the severity of disease expression. Ochola et al. (2014) observed low XW infections with a high occurrence of latent infections for inoculations using lower Xvm concentrations of 10^4^ cfu, whereas a high XW severity occurred at higher Xvm concentrations above 10^6^ cfu. In a recent study, a higher XW infection was also observed in enset (*Ensete ventricosum*; “false banana”) plants in which three leaves were inoculated compared with single leaves (Said et al. 2021).

### Xanthomonas vasicola pv. musacearum distribution within plant parts post inoculation

In the treatments in which two leaves had been inoculated, Xvm was only recovered in the pseudostem section next to the corm at 73 dpi compared to 43 days when all leaves were inoculated (Fig. 3A). A significantly higher (p =0.027) number of plants in which all leaves got inoculated (33%) had Xvm in the pseudostem section next to the corm compared with 17% for those in which only two leaves had been inoculated, further stressing the importance of the amount of Xvm inoculum in disease progression. In contrast to the sampled pseudostem sections, Xvm recovery in the mother plant corm tissues occurred much earlier (29 dpi) in both treatments, with no significant differences (p >0.05) in the recovery of Xvm from corm tissue samples between the two treatments (Fig. 3B). The time duration for Xvm to reach corm tissues in the current study is consistent with findings of Ocimati et al. (2013b) for vegetative stage inoculated plants. In addition to the delayed infections in mother plant corms, unexpectedly more mother plant corms (77-82%) had the Xvm bacteria compared to the pseudostem tissues. The lower and delayed recovery of Xvm in the pseudostem section close to the corm compared to the corm tissues is surprising. Xvm has been reported to occupy pockets of the vascular bundles (Tripathi et al. 2009; Blomme et al. 2017b) and could thus have been missed in the sampled pseudostem tissues assessed in the laboratory.

**Figure 3.**
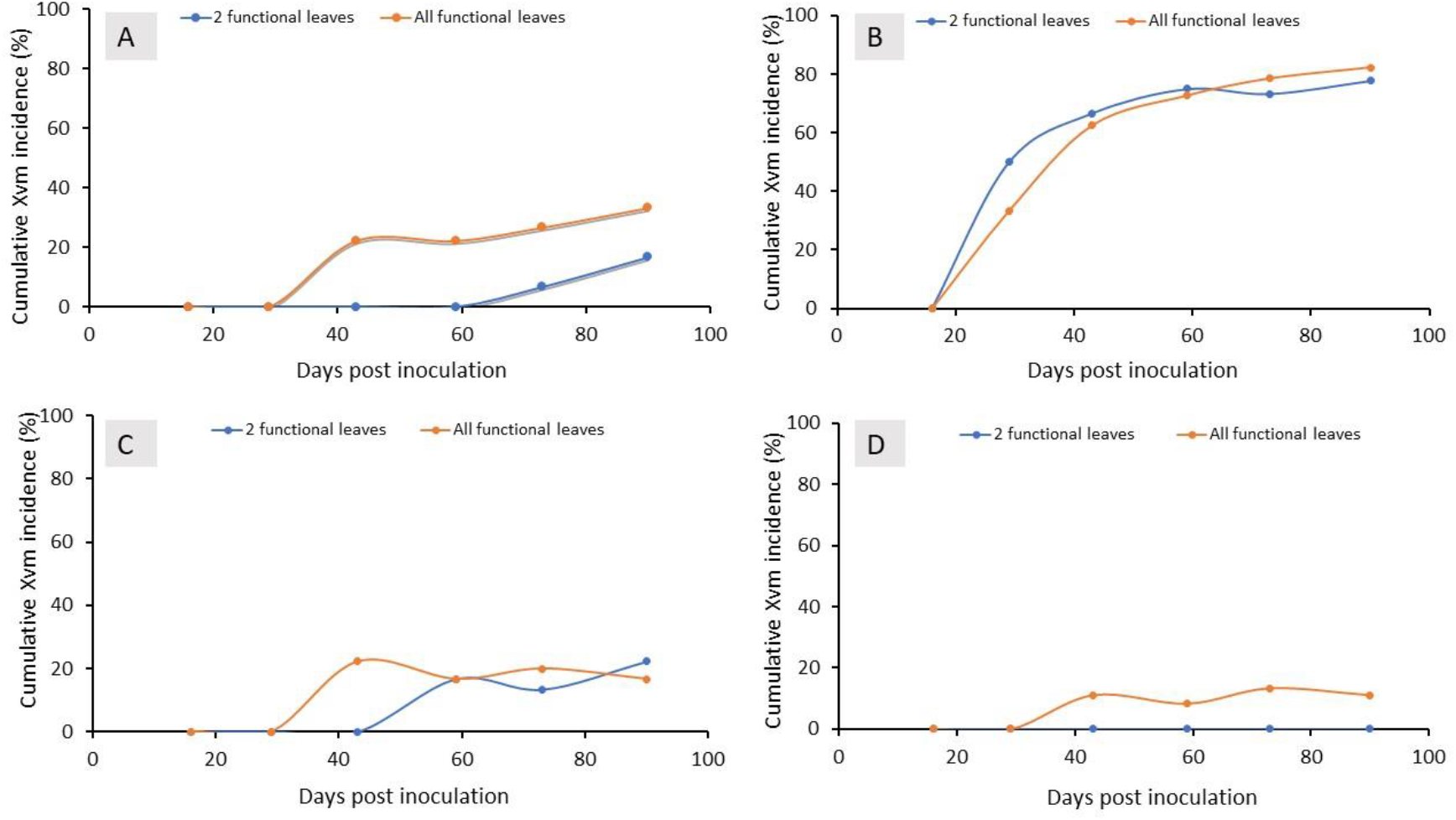
Cumulative *Xanthomonas vasicola* pv. *musacearum* (Xvm) incidences (%) in the A: mother plant pseudostem section next to corm tissue; B: corm tissues of the mother plant; C: sucker corm tissues; and D: in the cross section of the sucker pseudostem 10 cm from the sucker corm according to days post inoculation of mother plants with Xvm.

In the corms of the suckers, similarly Xvm was first recovered in the ‘all’ leaves treatment at 43 dpi compared to 59 dpi in the ‘two’ leaves treatments (Fig. 3C). Cumulatively, Xvm was only recovered from 22% of the corms. For the sucker pseudostems, Xvm was only retrieved from the ‘all leaves’ treatment and starting from 43 dpi (Fig. 3D).

No Xvm colonies were recovered from roots of both the mother plants and the attached suckers irrespective of the number of leaves inoculated and time from inoculation to sampling of plants. Ocimati et al. (2013b, 2011) also reported a lower colonization of banana root tissues.

### Xanthomonas vasicola pv. musacearum distribution within mother plant corm sections

The frequency of Xvm recovery in the lower, middle, and upper sections of the corm (Fig. 1B, C) tended to increase with the number of days post inoculation (Table 2). There was also a general tendency to have a lower Xvm incidence in the lower corm section relative to the middle and upper sections (Tables 2, Fig. 4). This could be explained by the fact that the upper corm section is directly attached to the leaf sheaths through which the inoculum gradually moves downwards. The trends in Xvm incidence between the two treatments where however not consistent (Tables 2, Fig. 4).

**Table 2.**
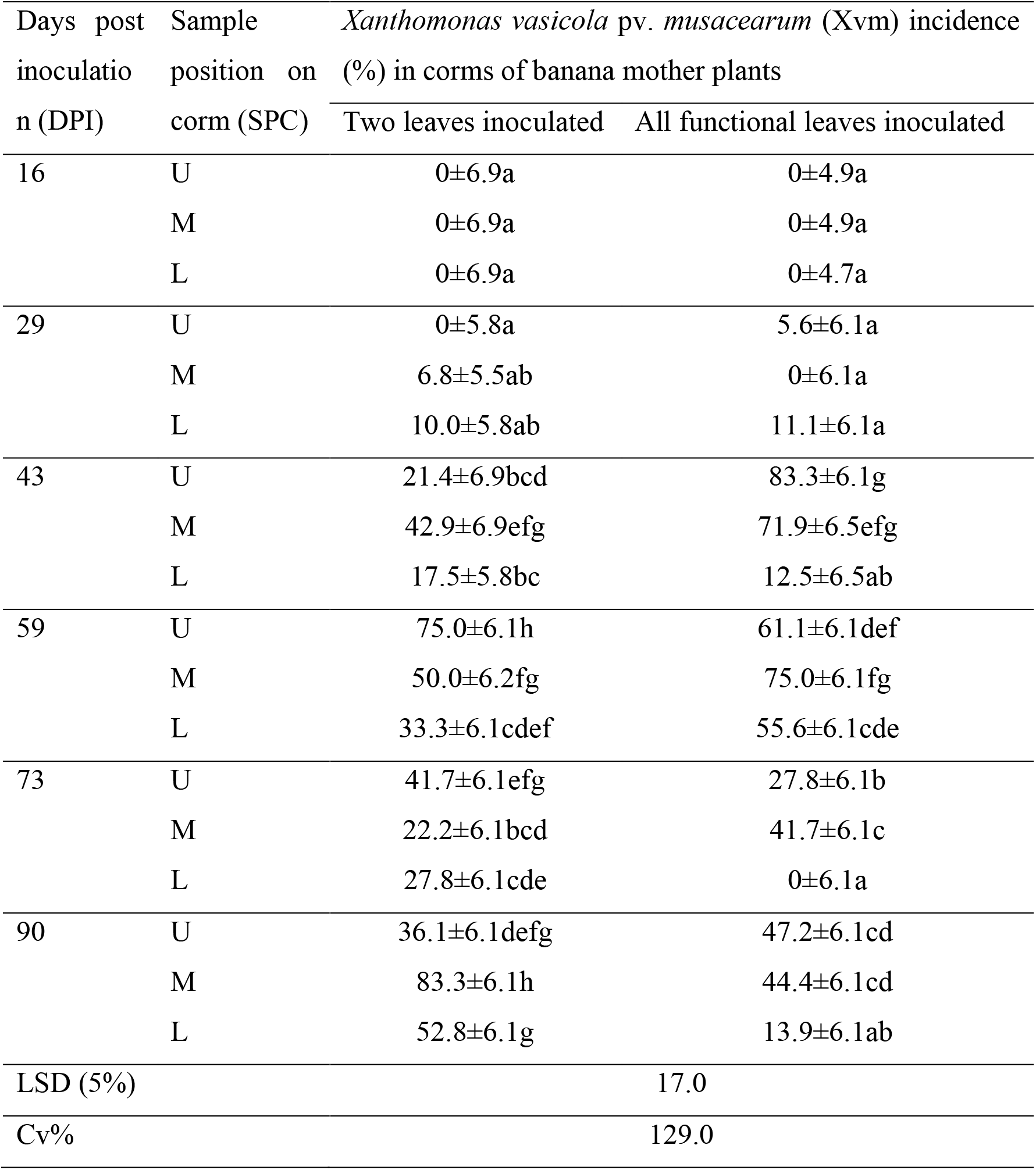
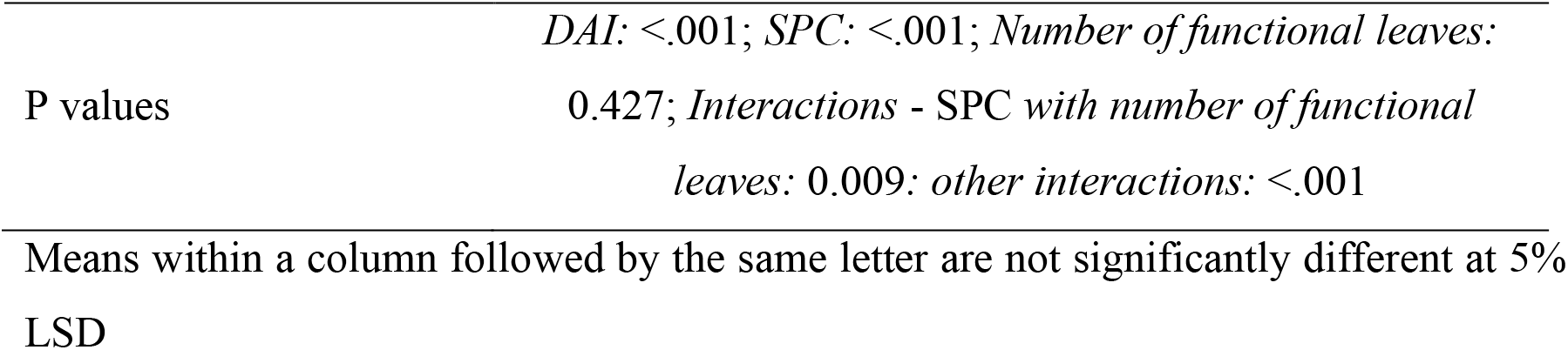
The incidence (%) of *Xanthomonas vasicola* pv. *musacearum* (Xvm) on the upper (U), middle (M) and lower (L) sections of corms of banana plants at 16, 29, 43, 59, 73 and 90 days after inoculation (DAI) following inoculation of two or all the functional leaves with a suspension of Xvm.

**Figure 4.**
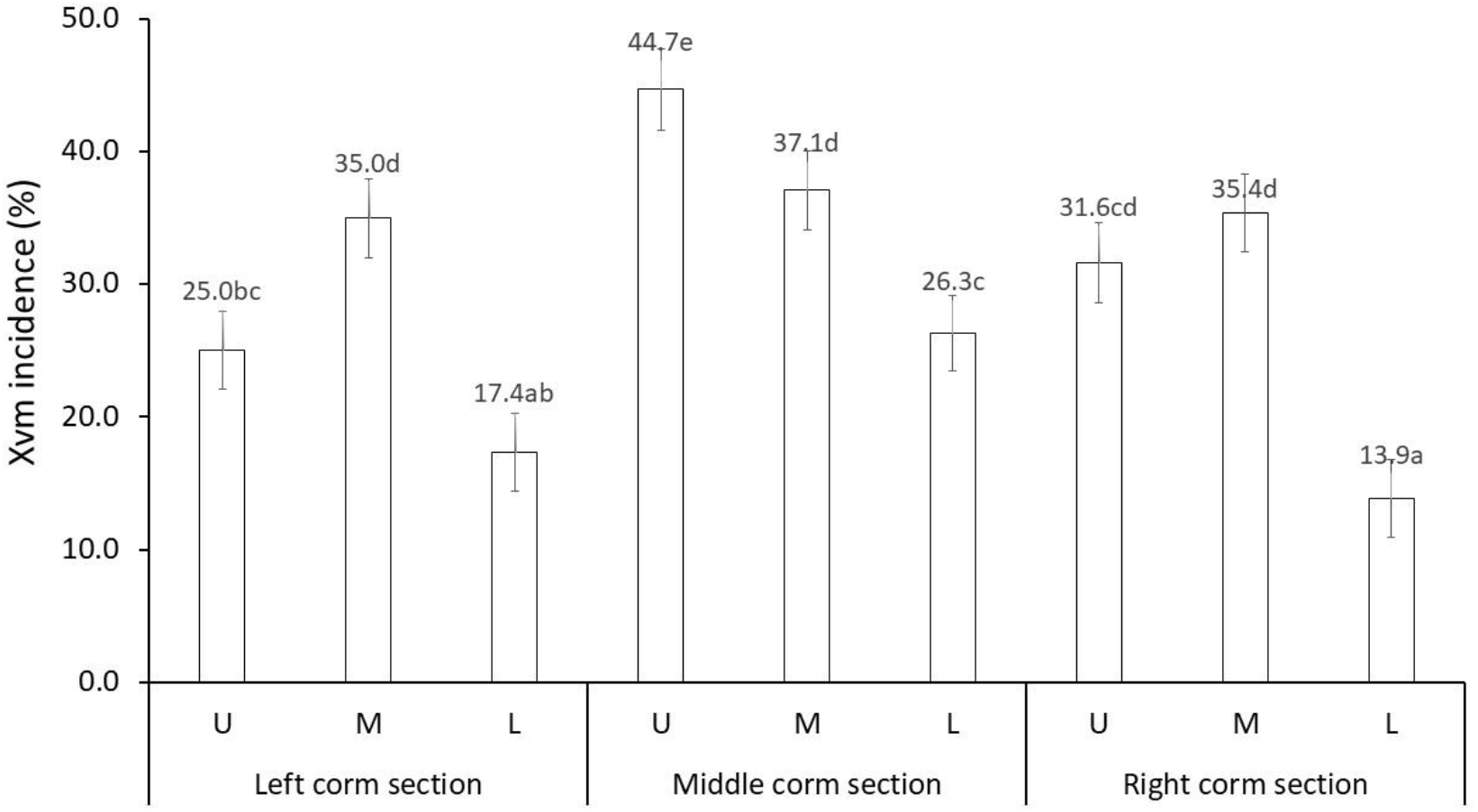
Comparison of the percentage incidence of *Xanthomonas vasicola* pv. *musacearum* (Xvm) in the different sections of the mother plant corms following inoculation of 6-month-old mother plants through all and two functional leaves. ‘U’, ‘M’ and ‘L’ respectively, denote upper, middle, and lower corm sections. Means followed by the same letter in are not significantly different at 5% LSD.

Higher Xvm recoveries also occurred for the middle corm zone made up of the central cylinder compared with the outer zones that comprised the cortex and layer of Mangin (Fig. 4). The upper middle corm section that is directly attached to the youngest leaf sheaths had the highest Xvm recoveries (Fig. 4) and was also observed to be significantly (p <0.01) softer than the other corm sections. The compactness of the tissues could have possibly also contributed to the observed distribution of Xvm in corm tissues. The upper corm sections were consistently softer than that of the middle and lower corm sections whereas the cortex was more compact than the central cylinder. The lower corm tissues on which most of the suckers are attached are the most compact corm tissues. The tissue hardness scores (kgf/cm^2^) for the central cylinder of vegetative stage plants were, respectively, 1.0, 1.7 and 2.0 for the upper, middle, and lower corm sections (Fig. 5A). Corm hardness readings for the cortex were consistently and significantly higher (p <0.01) than those for the central cylinder and were, respectively, 1.4, 2.3 and 2.8 for the upper, middle, and lower corm sections.

**Figure 5.**
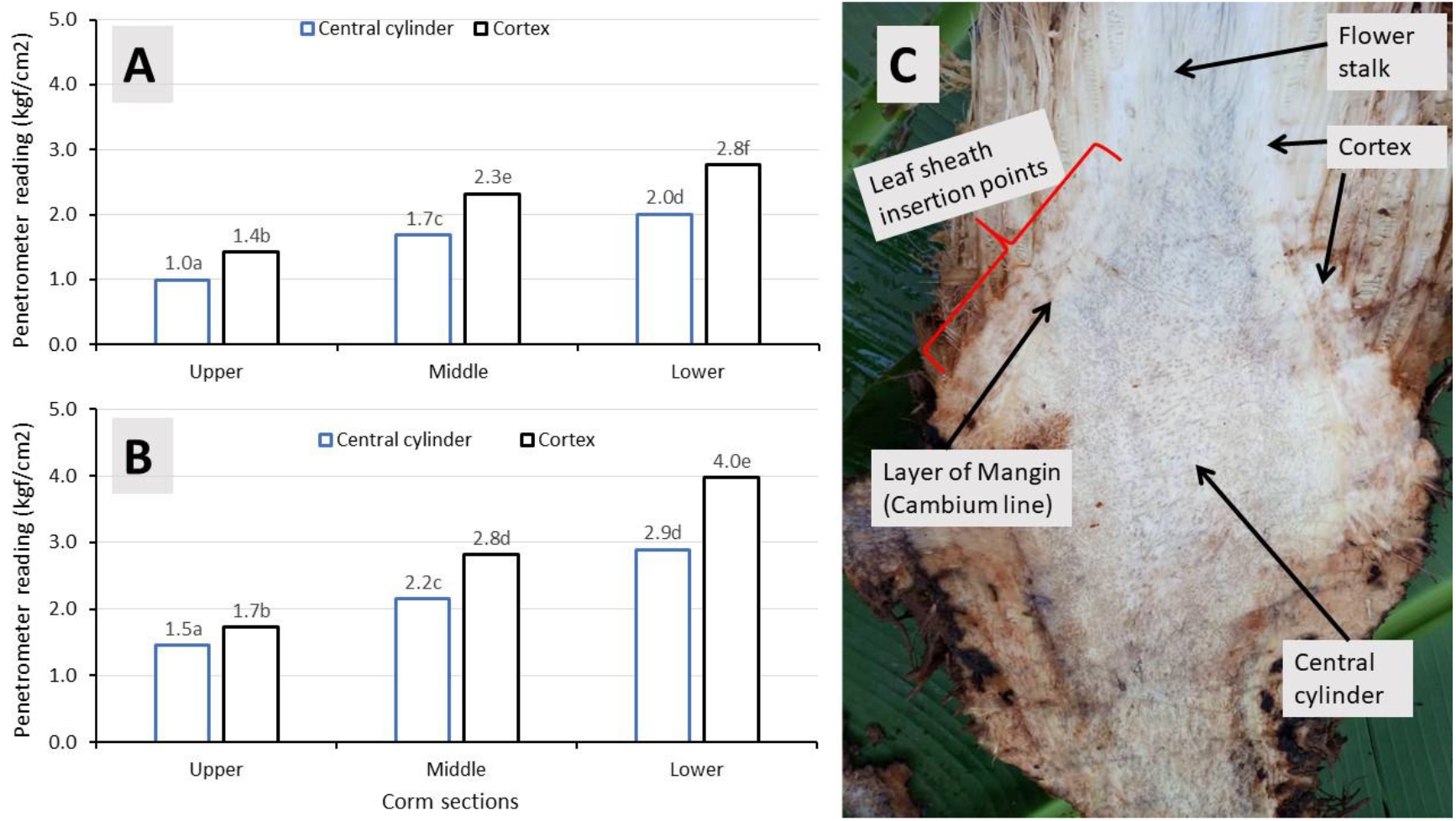
Corm hardness scores (kgf/cm^2^) for the upper, middle, and lower cortex and central cylinder sections of A) a vegetative stage mother plant and B) a flowering stage mother plant. C) A longitudinal section of the corm with an attached portion of the flower stalk (i.e., the real stem). Means followed by the same letter in (A) and (B) are not significantly different at 5% LSD.

The higher Xvm recoveries from the central cylinder of the corm (c.f. Fig. 4) contrast the findings of Ssekiwoko et al. (2006), who reported a higher recovery of bacteria in the layer of Mangin compared to the corm’s central cylinder or the cortex layer. According to Ssekiwoko et al. (2006), the structure of the cortex and central cylinder do not allow for a rapid Xvm spread. The layer of Mangin is made up of a mass of vascular bundles while the central cylinder and cortex are dominated by a mass of starchy parenchyma (Stover and Simmonds, 1987). The observed differences with the current study can be attributed to the differences in the points of entry of the bacteria. In the current study, Xvm was introduced through the leaves whereas through the floral parts or flower stalk (i.e., real stem) in Ssekiwoko et al. (2006). The flower stalk starts at the corm and continues all the way to the inflorescence or bunch (Fig 5C). In the current study, the corm sections of flowering stage plants were observed to be 21 to 50% more compact than the corm tissue sections of plants in the vegetative stage (Fig. 5B). The corm hardness scores (kgf/cm^2^) for the central cylinder of the flowering stage plants were, respectively, 1.5, 2.2 and 2.9 for the upper, middle, and lower corm sections while respectively, 1.7, 2.8 and 4.0 for the cortex. The current study did not explicitly separate the cortex from the layer of Mangin but observed the cortex tissue to be more compact than that of the central cylinder. Nevertheless, the observed differences in corm compactness could have also potentially affected Xvm spread and build up within them.

The logistic regression model showed Xvm incidence in the corm to be strongly (p <0.01) and positively influenced by the number of maiden suckers on the banana plants and the dpi (Table 3). The positive association between the number of maiden suckers and presence of Xvm in mother plant corm could be attributed to the higher demand for assimilates by the larger maiden suckers from the parent plants. The strong positive association between infections in the mother plant and infection in maiden suckers was reported by Ntamwira et al. (2019). Infections in medium to large-sized suckers causes frustration among farmers when applying the SDSR package (Blomme et al. 2019). Making this knowledge explicit to the farmers would reduce farmers’ frustration and minimise the risk of dis-adopting the control package. The increase in Xvm incidence in corm tissues with the dpi was expected and can be attributed to the increased build-up of inoculum in the corm tissues over time. This gives credence to the need to immediately remove symptomatic banana plants.

**Table 3.** Regression model of *Xanthomonas vasicola* pv. *musacearum* as the dependent variable with different independent variables.

| Variable | Estimate | Std error | t value | Pr (>\|t\|) |
| --- | --- | --- | --- | --- |
| (Intercept) | -0.1787 | 0.3160 | -0.566 | 0.5728 |
| Pseudostem girth at soil level | 0.0102 | 0.0060 | 1.701 | 0.0918 |
| Number of maiden suckers | 0.0622 | 0.0219 | 2.833 | 0.0054 |
| Height of mother plant corm | -0.0185 | 0.0079 | -2.325 | 0.0219 |
| Days post inoculation | 0.0027 | 0.0010 | 2.643 | 0.0094 |
| Number of functional leaves | -0.0473 | 0.0299 | -1.580 | 0.1169 |
| Mother plant height | -0.0027 | 0.0011 | -2.340 | 0.0211 |

Null deviance: 6.5847 on 117 degrees of freedom; Residual deviance: 5.5609 on 110 degrees of freedom

Xvm incidence within the corm tissues was also significantly (p <0.05) influenced by height of the mother plant corm and mother plant pseudostem height. The incidence of Xvm declined with increasing mother plant pseudostem and corm height (Table 3). The strong negative association (between Xvm incidence in the corm tissues with the height of the corm suggests that the further away the suckers are attached from the point of attachment of leaves and the apical meristem, the lower the risk of infection and vice versa. Most of the suckers were observed to be attached at the bottom of the corm tissues while the leaf sheaths through which the bacteria were introduced are attached to the upper and middle sections of the corm (c.f. Fig. 1A, B). This could partially explain the lower incidence of infections in suckers even when the mother plant or other attached plants in a mat are diseased. The negative association between plant height and Xvm incidence in corms can be attributed to the longer time duration Xvm needs to move through the pseudostems’ phloem vessels before reaching corm tissues in tall plants relative to the shorter ones. Ocimati et al. (2013b) also observed a weak positive correlation between plant height and time to symptom expression in banana plants.

These findings stress the importance of practices that reduce the amount of Xvm inoculum on banana mats and fields. Ntamwira et al. (2019) observed a faster recovery of mats through SDSR when diseased plants are removed as soon as symptoms are observed. In these plants, a lower amount of ooze was observed to build up compared to when the SDSR application was delayed for multiple weeks. Said et al. (2021) also demonstrated early removal of visibly diseased outer leaf sheaths on infected enset plants to reduce leaf symptom incidence and increase plant recovery. Ssekiwoko et al. (2010) and Ocimati et al. (2013a) also reported that entire banana mats could be saved if florally infected banana plants showing early male bud infections were immediately cut. Even under worst case scenarios, with high initial plant incidence levels and where farmers in frustration cut all banana pseudostems in their infected fields with a single tool, thus potentially spreading the disease to all mats, emergence of healthy-looking shoots and recovery of entire fields have been observed (Blomme et al. 2017a). Thus, even when Xvm bacteria reach the corm tissues of an infected plant, disease progression to the physically attached shoots is incomplete/hampered. This finding could partly explain the current success achieved through singly removing diseased banana plants as a method for managing XW disease in banana (Blomme et al. 2014, 2017a, 2019).

This study was conducted in a single location as such effects of variation in soil and environmental conditions could not be ascertained. The study was also limited in terms of the coverage of banana cultivars or genome groups, given potential differences could have arisen between cultivars or genome groups. Nevertheless, the success of SDSR across the different regions of East and Central Africa (Blomme et al. 2014, 2017a, 2019, 2021; Kubiriba et al. 2012; Ocimati et al. 2013a, 2015; Ntamwira et al. 2019), suggests that the impact of these variables could be minimal.

## Conclusion

This study suggests the possibility of a combination of multiple factors including the physiology of the banana plant in influencing Xvm spread within the banana mat. The chance of an attached sucker to get infected from the mother plant is higher when it is taller. Making this knowledge explicit to farmers as part of the extension package on the use of SDSR is crucial and will reduce the risk of dis-adopting the control practice when farmers observe new and especially large diseased plants in treated mats. The fact that most lateral shoots are connected to the bottom part/lower half of the mother plant corms delayed or prevented their infection and the fact that the dpi and amount of inoculum played a role in infection suggests that timely removal of diseased plants will minimize infections in the attached suckers. This study thus confirms the need for the immediate removal of diseased stems on a mat in order to reduce Xvm inoculum levels in a timely manner. This knowledge will enhance the confidence of extension personnel and farmers in the SDSR control package. Studies to understand the possible role of different corm tissues in Xvm movement, for a range of Musa cultivars, is recommended.

## Author contributions

Conceptualization, Walter Ocimati and Guy Blomme; Data curation, Walter Ocimati; Formal analysis, Walter Ocimati; Investigation, Walter Ocimati and Anthony Tazuba; Methodology, Walter Ocimati and Guy Blomme; Project administration, Guy Blomme; Resources, Guy Blomme; Supervision, Guy Blomme; Visualization, Walter Ocimati; Writing – original draft, Walter Ocimati and Anthony Tazuba; Writing – review & editing, Walter Ocimati, Anthony Tazuba and Guy Blomme.

## Acknowledgments

The authors are grateful for the support of the Directorate General for Development (DGD-Belgium) through the CIALCA project, the CGIAR Research Program on Roots, Tubers and Bananas (RTB) and the CGIAR Trust Fund contributors. the CGIAR Research Program on Roots, Tubers and Bananas. We also acknowledge Ms. Florence Nakamanya, Ms. Dorcus Nassazi and Leniru Salome for their assistance with laboratory work.

## Data availability statement

The raw data supporting the conclusions of this manuscript will be made available by the authors, without undue reservation, to any qualified researcher.

